# Duplicated zebrafish (*Danio rerio*) inositol phosphatases *inpp5ka* and *inpp5kb* diverged in expression pattern and function

**DOI:** 10.1101/2022.08.31.506059

**Authors:** Dhyanam Shukla, Brian M. Gural, Edmund S. Cauley, Llion E. Roberts, Brittany F. Karas, Luca Cavallo, Luka Turkalj, Sally A. Moody, Laura E. Swan, M. Chiara Manzini

**Affiliations:** Department of Neuroscience and Cell Biology and Child Health Institute of New Jersey, Rutgers Robert Wood Johnson Medical School, New Brunswick, NJ, USA; Department of Biochemistry and Molecular Medicine, School of Medicine and Health Sciences, The George Washington University, Washington, DC, USA; Institute of Systems, Molecular and Integrative Biology, University of Liverpool, Liverpool, UK; Department of Anatomy and Cell Biology, School of Medicine and Health Sciences, The George Washington University, Washington, DC, USA

**Keywords:** inositol phosphatase, INPP5K, zebrafish, gene duplication

## Abstract

One hurdle in the development of zebrafish models of human disease is the presence of multiple zebrafish orthologs resulting from whole genome duplication in teleosts. Mutations in Inositol polyphosphate 5-phosphatase K (*INPP5K*) lead to a syndrome characterized by variable presentation of intellectual disability, brain abnormalities, cataracts, muscle disease, and short stature. INPP5K is a phosphatase acting at position 5 of phosphoinositides to control their homeostasis and is involved in insulin signaling, cytoskeletal regulation, and protein trafficking. Previously, our group and others have replicated the human phenotypes in zebrafish knockdown models by targeting both *INPP5K* orthologs *inpp5ka* and *inpp5kb*. Here, we show that *inpp5ka* is the more closely related orthologue to human *INPP5K*. While both *inpp5ka* and *inpp5kb* mRNA expression levels follow a similar trend in the developing head, eyes, and tail, *inpp5ka* is much more abundantly expressed in these tissues than *inpp5kb. In situ* hybridization revealed a similar trend, also showing unique localization of *inpp5kb* in the pineal gland indicating different transcriptional regulation. We also found that *inpp5kb* has lost its catalytic activity against its preferred substrate, PtdIns(4,5)P_2_. Since most human mutations are missense changes disrupting phosphatase activity, we propose that loss of *inpp5ka* alone can be targeted to recapitulate the human presentation. In addition, we show that the function of *inpp5kb* has diverged from *inpp5ka* and may play a novel role in the zebrafish.

## Introduction

Inositol polyphosphate 5-phosphatase K (INPP5K [MIM:607875]) is a highly conserved phosphatase that participates in the regulation of phosphoinositide (PI) signaling. Also referred to as skeletal muscle and kidney-enriched inositol phosphatase (*SKIP*), *INPP5K* is highly expressed in the brain, eyes, and muscles during development and adulthood (Gurung et al., 2003; Ijuin et al., 2000). In humans, homozygous or compound heterozygous mutations in *INPP5K* have been causally linked to a form of congenital muscular dystrophy with cataracts and intellectual disability (MIM: 617404) also associated with short stature, and microcephaly with considerable variability in the age of onset and early presentations (D’Amico et al., 2020; Hathazi et al., 2021; Osborn et al., 2017; Wiessner et al., 2017; Yousaf et al., 2017). Similarities have been noted with Marinesco-Sjögren syndrome (MIM: 248800), a form of myopathy also associated with congenital cataracts, short stature, and cerebellar ataxia (Krieger et al., 2013; Senderek et al., 2005).

PIs are a category of lipid molecules that play crucial roles in signal transduction, ion channel regulation, cellular migration, membrane trafficking, vesicle transport, and many other processes (Balla, 2013; Paolo and Camilli, 2006; Raghu et al., 2019). The seven unique members of this group are distinguished by their patterns of phosphorylation of the phosphorylated inositol head (PtdIns), which can occur at one or more of three positions (−3, −4, or −5). Production of PIs is regulated by an array of kinases and phosphatases (Balla, 2013). INPP5K hydrolyzes the D-5 position of the inositol ring in both PtdIns(4,5)P_2_ and PtdIns(3,4,5)P_3_, with highest activity for PtdIns(4,5)P_2_ (Davies et al., 2015; Ijuin et al., 2000; Vandeput et al., 2006). INPP5K is largely localized to the endoplasmic reticulum (ER) (Dong et al., 2018; Gurung et al., 2003) but can translocate to membrane ruffles as part of a complex with the glucose regulated protein GRP78/BiP to negatively regulate insulin receptor signaling via phosphatidylinositol-3-kinase (PI3K) (Ijuin and Takenawa, 2003; Ijuin et al., 2015, 2016a, 2016b).

Multiple zebrafish (*Danio rerio*) models of *INPP5K* loss of function have been generated using morpholino oligonucleotides (MOs) targeting both paralogs, *inpp5ka* and *inpp5kb* (Hathazi et al., 2021; Osborn et al., 2017; Wiessner et al., 2017). However, when the genes were targeted independently, knockdown of *inpp5ka* was sufficient to yield phenotypes typical of neurological and muscular disorders, such as microphthalmia, microcephaly, shortened body, reduced touch-evoked motility and myopathy. In contrast, *inpp5kb* MOs produced a mild phenotype in a small subset of morphants (Osborn et al., 2017). In addition, we found *inpp5ka* expression to be 16-fold higher than *inpp5kb* in zebrafish embryos at 2 days post fertilization (dpf) (Osborn et al., 2017). These findings suggested that *inpp5ka* may be the most conserved human paralog and *inpp5kb* function may have diverged.

Due to a genome duplication event in teleost fish, about 30% of zebrafish genes have a paralog (Howe et al., 2013), but duplicated genes often acquire differential expression and function (Postlethwait et al., 1998; Ravi and Venkatesh, 2018). In this study we sought to better characterize expression patterns and function of *inpp5ka* and *inpp5kb* to understand whether they diverged and support the development of better models of *INPP5K* mutations in humans. We show that both *inpp5ka* and *inpp5kb* have a dynamic developmental expression in the eyes, head, and tail, also finding that despite being expressed at lower levels, *inpp5kb* is specifically enriched in the pineal gland. Inpp5kb lost the majority of its phosphatase activity for PtdIns(4,5)P_2_ which is the preferred substrate for INPP5K (Ijuin et al., 2000). Together, these data indicate that *inpp5ka* is the closest ortholog to *INPP5K* and suggest a unique role for *inpp5kb* within the zebrafish.

## Methods

### Animal Care

Maintenance and husbandry of zebrafish (*Danio rerio*) breeders and larvae were performed following protocols approved by the Institutional Animal Care and Use Committee of the George Washington University and Rutgers University. All animals were from the AB background.

### Protein alignments

QIAGEN CLC Sequence Viewer 8 was used to align the sequences for all transcripts. Percent identities between the human INPP5K (NP_057616.2), zebrafish Inpp5ka (NP_001082962.2), and Inpp5kb (XP_021335020.1) were calculated using EMBOSS Needle (Madeira et al., 2022).

### Quantitative PCR (qPCR) analysis

Samples were collected at 1, 2, 3, 4 and 5 dpf. Whole zebrafish embryos and larvae or micro-dissected tissue from eyes, head, and tails were pooled and RNA was extracted using the ReliaPrep RNA Miniprep System kit (Promega, Madison, WI). RNA was treated with DNase I (New England Biolabs, Ipswich, MA) and complementary DNA (cDNA) was synthesized using the iScript cDNA Synthesis kit (Bio-Rad, Hercules, CA). 600 ng of cDNA per sample were analyzed via qPCR using the SsoFast EvaGreen Supermix (Bio-Rad, Hercules, CA) on a Bio-Rad CFX384 Touch Real Time PCR System. All reactions were run with 3 technical replicates and repeated on at least 3 biological replicates. Sequences for custom primers for *inpp5ka* and *inpp5kb* and housekeeping controls elongation factor 1 alpha *eef1a* and riboprotein L13 *rpl13* are available upon request.

### Whole-mount in situ hybridization (ISH)

Full length *inpp5ka* (NM_001089493.1) and *inpp5kb* (XM_021479345.1) cDNAs were cloned into the pCS2+ plasmid (Addgene, Watertown, MA). Digoxygenin-labeled sense and antisense probes were synthesized from the linearized plasmids using the DIG RNA Labeling Kit (SP6/T7) (Roche/MilliporeSigma, Burlington, MA). Whole-mount ISH was performed as previously described (Yan et al., 2009).

### Phosphatase assay

Full length *inpp5ka* (NM_001089493.1) and *inpp5kb* (XM_021479345.1) cDNAs were generated by gene synthesis and cloned into the pGEX-1 to generate GST-fusion proteins (Genewiz/Azenta Life Sciences, South Plainfield, NJ). GST-human INPP5K and GST were used as positive and negative controls respectively (Weissner et al., 2017). Constructs were transformed into BL21 DE3 pLysS, induced with 100uM IPTG overnight and harvested by centrifugation. Cells were lysed in assay buffer (50 mM Tris-HCl [pH 7.5], 150 mM NaCl, 10 mM MgCl_2_) plus 1% Triton X-100, EDTA-free protease inhibitors (Roche Diagnostics) and turbonuclease (Sigma). GST fusion proteins were affinity purified over Gluthione Sepharose 4B (GE Healthcare). After extensive washing, aliquots of beads were run on Coomassie gels to determine the abundance of full-length fusion proteins. Beads bearing equal amounts of fusion proteins were incubated in assay buffer containing 135 μM PtdIns(4,5)P_2_diC8 or PtdIns(3,4,5)P_3_diC8, including control wells with no enzyme or no substrate lipid, and incubated for 1h at 37C. Free phosphate was measured using the Malachite Green assay kit (Echelon Biosciences). Results of three independent experiments were presented as mean ± standard deviation. To minimize variability between purifications, all constructs were freshly prepared and purified in parallel for each experiment.

## Results

### Zebrafish and human INPP5K protein alignments

To determine whether *inpp5ka* and *inpp5kb* lead to functionally divergent proteins, we first analyzed their protein sequence. Protein sequence alignment of INPP5K (NP_057616.2), Inpp5ka (NP_001082962.2), and Inpp5kb (XP_021335020.1) (**Fig. 1**) revealed 42.6% and 38.5% identity between the human orthologue and Inpp5ka and Inpp5kb respectively, while the zebrafish proteins showed 56% identity with each other. Inpp5kb has an additional 48 amino acid N-terminal sequence that was not present in either Inpp5ka or INPP5K.

**Figure 1.**
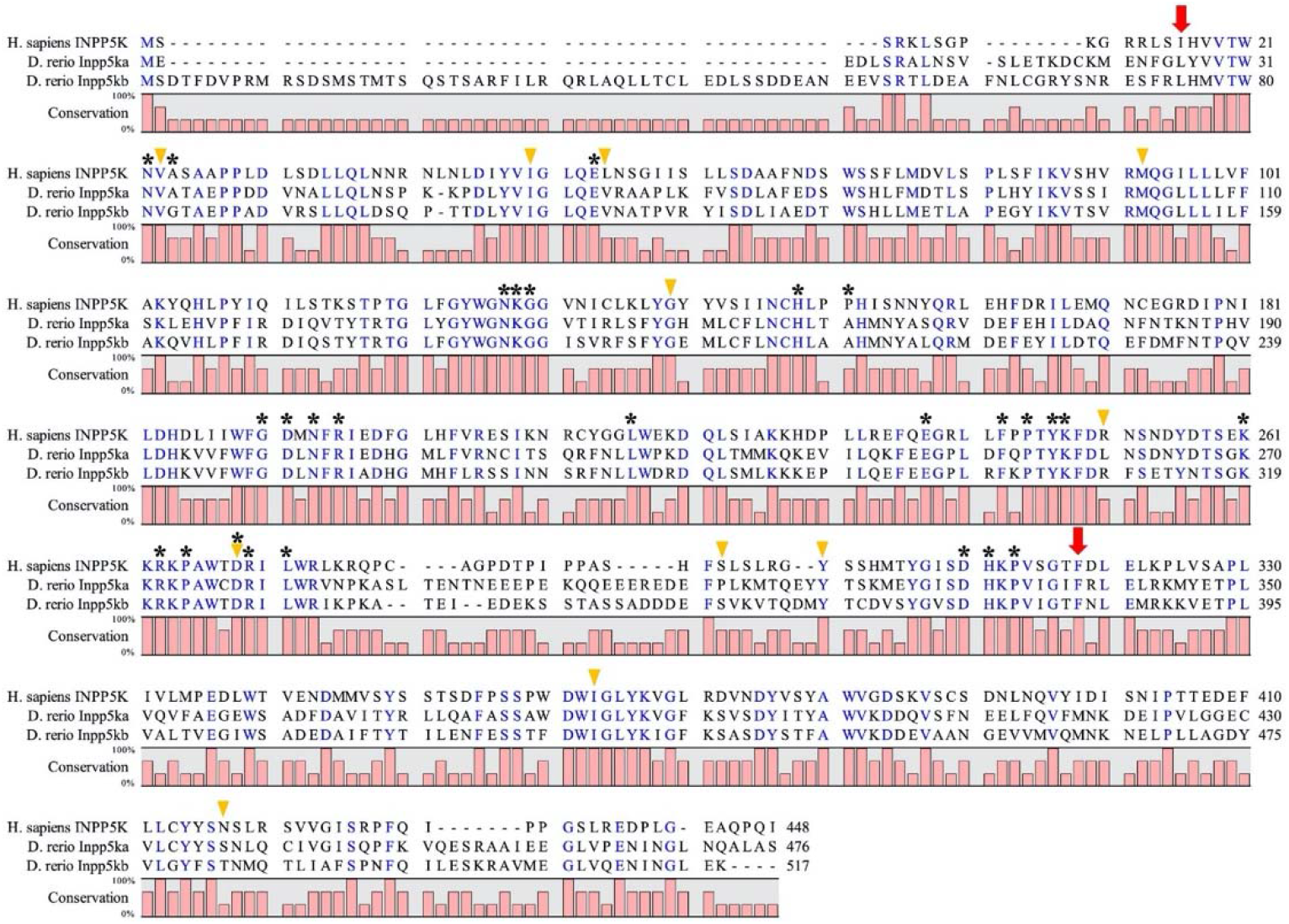
Protein alignment of INPP5K, Inpp5ka and Inpp5kb highlighting conserved amino acids required for phosphatase activity. The start and end of the catalytic domain in the human protein are marked with arrows. Amino acids required for phosphatase activity have been denoted with asterisks (*). Arrowheads indicate residues that are altered by missense variants in humans.

### Divergent expression and localization of INPP5K orthologs in zebrafish larva

Analysis of *inpp5ka* and *inpp5kb* mRNA obtained from whole zebrafish embryos had shown higher expression of *inpp5ka* (Osborn et al., 2017). We used qPCR to define expression patterns throughout the first five days of development. We found that *inpp5ka* (NM_001089493.1) was consistently expressed much more abundantly than *inpp5kb* (XM_021479345.1) (**Fig. 2A**). The developmental expression trend was similar for *inpp5ka* and *inpp5kb*. Both genes had relatively low levels of expression at 1- and 2-dpf, but gene expression peaked at 4 dpf, where we saw a 3.2-fold difference between the two paralogs. Gene expression started decreasing in the following day (**Fig. 2A**, Fold change relative to 1dpf *inpp5kb. inpp5ka*: 2dpf 11.1±1.5, 3dpf 51.9±0.4, 4dpf 112.1±3.6, 5dpf 84.9±5.7; *inpp5kb*: 2dpf 2.1±0.1, 3dpf 9.0±2.3, 4dpf 35.5±3.9, 5dpf 23.6±2.5. p>0.0001 at 3, 4, and 5 dpf).

**Figure 2.**
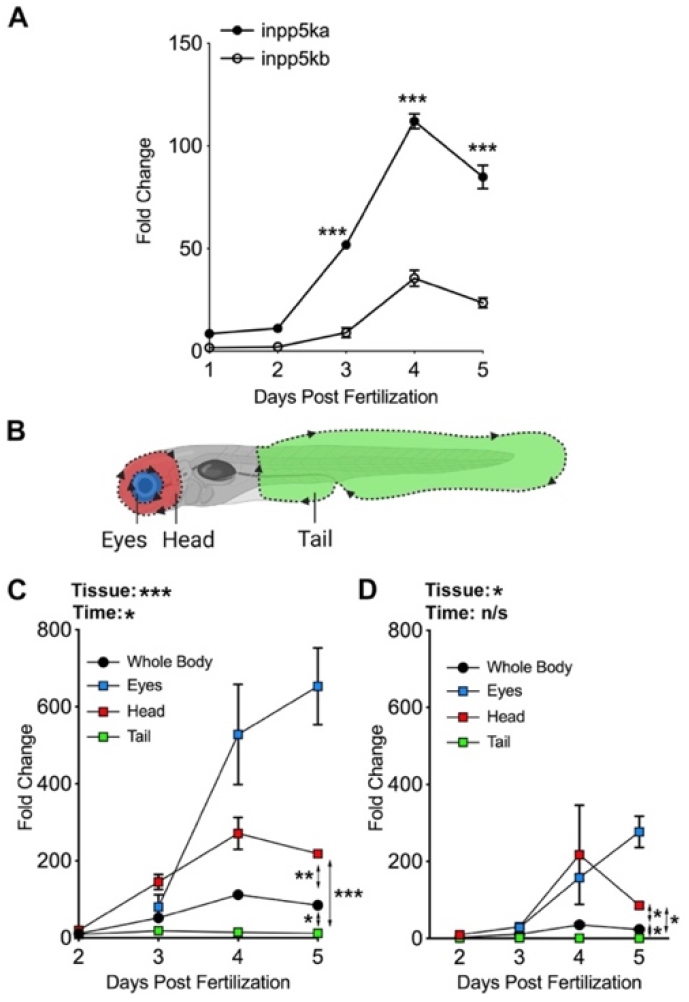
*inpp5ka* and *inpp5kb* mRNAs differ in expression in zebrafish larvae. **A.** Gene expression determined by qPCR. *inpp5ka* is more highly expressed in whole body lysates through 5 dpf. **B.** Larval tissues were excised from the eye, head, and tail for localized gene expression analysis. **C-D.** Expression for both *inpp5ka* (**C**) and *inpp5kb* (**D**) is low in the tail and increases in the eyes and brain. By 5 dpf, both are most highly expressed in the eyes. Values are averages ± SEM. * p < 0.05, ** p < 0.01, *** p < 0.001

Loss of *INPP5K* in humans affects the muscle, brain, and eyes and knockdown of *inpp5ka* in zebrafish larvae resulted in morphological abnormalities in the eyes and skeletal muscle (Hathazi et al., 2021; Osborn et al., 2017; Wiessner et al., 2017). We dissected the heads, eyes, and tails of developing larvae for tissue-specific expression analysis (**Fig. 2B**). This revealed that, while *inpp5ka* was consistently expressed at higher levels, both paralogs exhibit the greatest expression in the eyes and head. *inpp5ka* and *inpp5kb* showed 4.7- and 4.4-fold higher expression in the eyes respectively when compared to the whole-body at 4 dpf. The head revealed a 2.4-fold difference in *inpp5ka* and a 6.1-fold difference in *inpp5kb* compared to the whole-body expression (**Fig. 2C-D**, Fold change relative to 1dpf *inpp5kb. inpp5ka:* 3dpf head 145.2±19.3, eyes 80.4±31.6; 4dpf head 271.2±41.5, eyes 528.1±129.8; 5dpf 218.5±2.7, eyes 653.1±99.6. *inpp5kb:* 3dpf head 30.4±3.4, eyes 28.6±4.4; 4dpf head 217.3±128.8, eyes 157.7±8.4; 5dpf head 85.8±7.9, eyes 276.9±40.6).

To determine whether the expression patterns were consistent, we conducted *in-situ* hybridization on whole-mount larvae at 3 dpf. *inpp5ka* antisense probing reflected the results of our previous gene expression assays. *inpp5ka* mRNA was most abundant in the head and eyes, with lower expression in the tail (**Fig. 3A-C**). As expected, *inpp5kb* antisense targeting revealed moderate expression throughout the head and eyes (**Fig. 3D-E**). However, in contrast with *inpp5ka, inpp5kb* was abundantly expressed in the pineal gland (**Fig. 3F**), a neuroendocrine organ which responds to light and plays a role in circadian rhythm (Cahill, 1996; Livne et al., 2016; Vatine et al., 2011). These findings indicate that in addition to lower expression, *inpp5kb* also diverged in its expression pattern.

**Figure 3.**
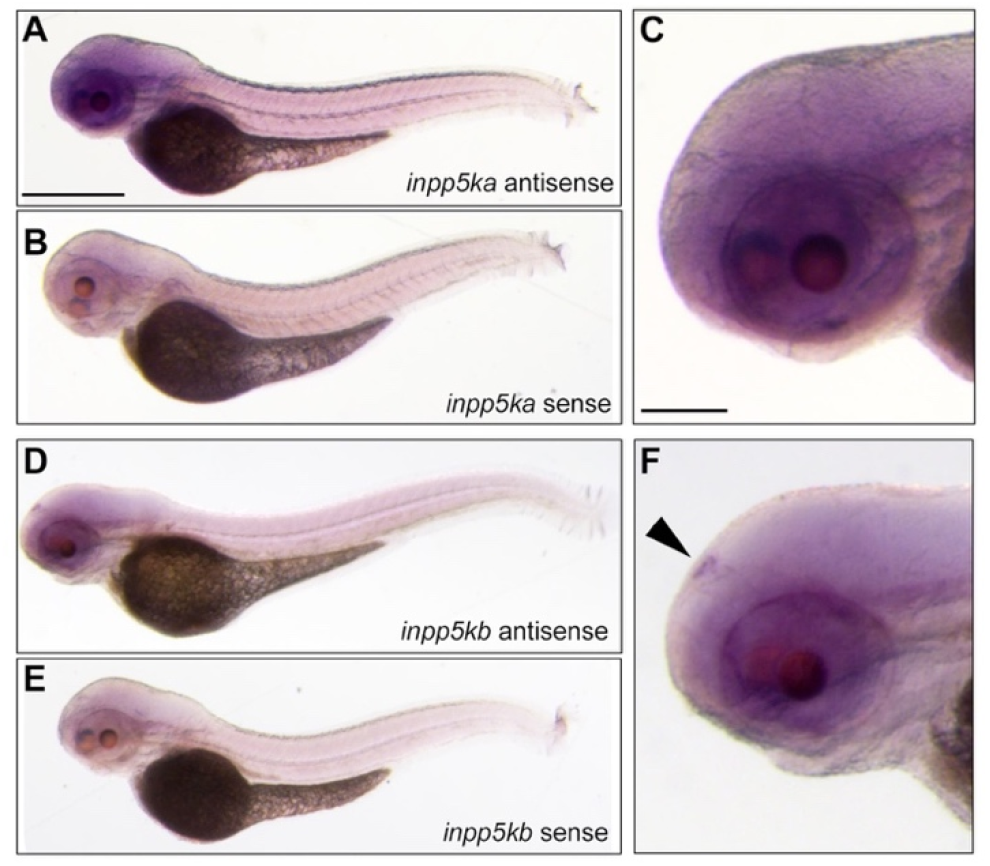
*inpp5ka* and *inpp5kb* mRNAs differ in localization in zebrafish larvae. **A-C.** *In situ* hybridization in 3 dpf larvae shows that *inpp5ka* mRNA is highly expressed throughout the head and eyes. Scale bars: 500μm in A, 100μm in C. **D-F**. *inpp5kb* expression is concentrated to the pineal gland. The pineal gland is indicated by the black arrow.

### Divergence in phosphatase activity of human and zebrafish orthologs of INPP5K

To evaluate the preservation of the PI phosphatase activity in the zebrafish isoforms, we conducted a malachite phosphatase assay to examine the activity of INPP5K and the two zebrafish Inpp5k isoforms against the preferred substrate PtdIns(4,5)P_2_ (**Fig. 4A)**. We found that zebrafish Inpp5ka and human INPP5K were both highly active against PtsIns(4,5)P_2_. This activity was specific, as illustrated by the lack of phosphatase activity against PIP_3_. However, compared to Inpp5ka, Inpp5kb was nearly inactive against PtdIns(4,5)P_2_. Inpp5ka yielded 409 pmol of free phosphate vs 20pmol for Inpp5kb, indicating that Inpp5ka had a 20-fold higher activity compared to Inpp5kb (**Fig. 4B)**.

**Figure 4.**
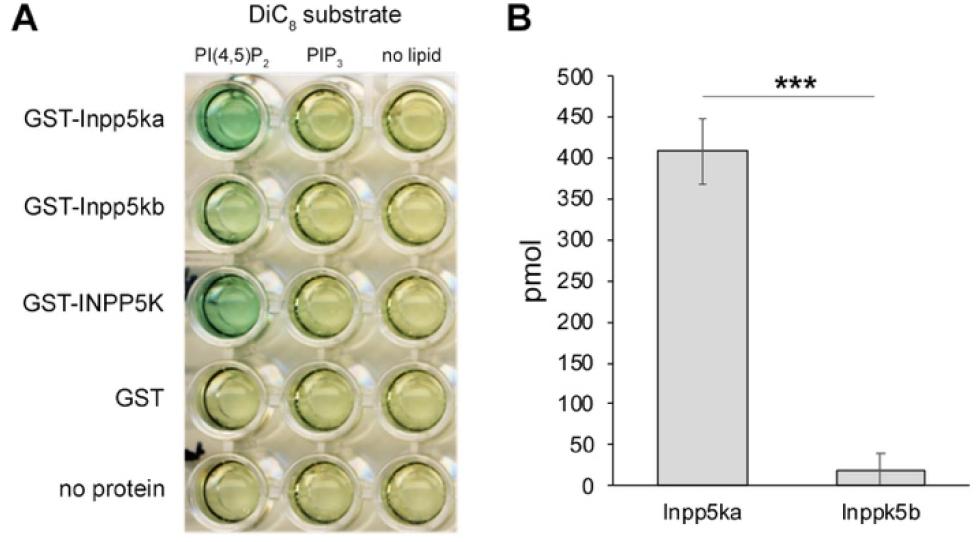
Inpp5ka and Inpp5kb exhibit different phosphatase activity. A. Phosphatase activity of human INPP5K, Inpp5ka and Inpp5kb in malachite assay. Human INPP5K and Inpp5ka demonstrate high activity for the PI(4,5)P_2_ substrate. PIP_3_ did not elicit activity from any isoform. **B.** Inpp5ka is more significantly active against diC8PI(4,5)P_2_ compared to Inpp5kb. Values are averages ± SEM. *** p < 0.001 following t-test.

The INPP5K protein is primarily composed of a 5-phosphatase domain between amino acids 16-318 and a SKITCH domain between amino acids 321-448. Most found mutations in humans are missense and have been shown to reduce or ablate phosphatase activity (Osborn et al., 2017; Wiessner et al., 2017). We wondered whether the loss in activity in Inpp5kb could be caused by changes in amino acids identified to be critical for the catalytic activity of INPP5K. Basing this analysis on the available crystal structures of other Type II inositol phosphate 5-phosphatases, *INPP5B* and *SYNJ1* (Paesmans et al., 2020; Trésaugues et al., 2014), we found that all sites were conserved in Inpp5ka and Inpp5kb and there were no major changes that could explain differences in activity (asterisks in **Fig. 1**). We also assessed whether residues known to be affected by pathogenic variants in patients were conserved in Inpp5kb, and these amino acids were all maintained (arrowheads in **Fig. 1**) (D’Amico et al., 2020; Osborn et al., 2017; Hathazi et al., 2021, Wiessner et al., 2017; Yousaf et al., 2017). Thus, possible changes on known residues do not explain the difference in function between Inpp5ka and Inpp5kb.

## Discussion

*INPP5K* mutations in humans cause a distinct neurodevelopmental syndrome with variable presentation of intellectual disability, cataracts, short stature, and muscle disease (D’Amico et al., 2020; Osborn et al., 2017; Wiessner et al., 2017; Yousaf et al., 2017). Multiple zebrafish models have been developed to study *inpp5k* function using morpholino oligonucleotides either blocking translation or knocking down mRNA expression (Hathazi et al., 2021; Osborn et al., 2017; Wiessner et al., 2017). However, the presence of duplicated *inpp5k* genes, *inpp5ka* and *inpp5kb*, in zebrafish complicates the development of both candidate loss- and gain-of-function mutations since both zebrafish orthologs may need to be targeted. Initial functional data from our previous studies had shown that *inpp5ka* knockdown alone was sufficient to replicate the findings in the double gene knockdown (Osborn et al., 2017). In this study, we show that *inpp5ka* and *inpp5kb* have diverged in expression levels, patterns and function following teleost whole genome duplication (WGD). *inpp5ka*, rather than *inpp5kb*, maintains a higher sequence identity to human *INPP5K*, suggesting that genetic removal of this gene may be sufficient to recapitulate the human mutation.

Polyploidization by WGD is a significant driver of evolution (Postlethwait et al., 1998; Sémon and Wolfe, 2007). During the period of re-diploidization that follows a WGD event, most redundant genes are eliminated via genomic rearrangements and mutations causing one duplicated copy to become a pseudogene. However, a duplicated gene may be preserved and gradually diverge in expression patterns and/or function during evolution leading to gene adaptation through sub-functionalization or neo-functionalization of one of the duplicated genes (Kassahn et al., 2009; Sémon and Wolfe, 2007). While *inpp5ka* is broadly and highly expressed throughout the zebrafish larvae, *inpp5kb* is significantly less expressed. Additionally, we found that Inpp5kb exhibits minimal phosphatase activity against the traditional substrate of INPP5K, PtdIns(4,5)P_2_ (Ijuin et al., 2000; Vandeput et al., 2006). While both paralogs are most abundant in the head, the visualization of expression achieved by *in-situ* hybridization reveals that the expression of *inpp5kb* is specifically enriched in the pineal gland. The pineal gland is thought to be the master regulator for circadian rhythm in vertebrates. Melatonin is the key circadian hormone secreted by the pineal gland in zebrafish (Cahill, 1996) and is thought to play a role in locomotor activity (Livne et al., 2016), as well in the timing of reproduction and feeding (Piccinetti et al., 2013). It will be interesting in the future to determine whether Inpp5kb is involved in pineal functions independently of its phosphatase activity.

In humans, much of the pathology resulting from mutations within *INPP5K* have been attributed to the dysregulation of phosphoinositide homeostasis (Hathazi et al., 2021; McGrath et al., 2020; Osborn et al., 2017; Wiessner et al., 2017). Most known mutations in INPP5K are missense variants occurring in the catalytic phosphatase domain reducing or ablating conversion of PtdIns(4,5)P_2_ to PtdIns(4)P (Osborn et al., 2017; Wiessner et al., 2017). In the muscle, INPP5K is involved in insulin signaling through the PI3K/Akt/mTOR pathway (Ijuin and Takenawa, 2015; Ijuin et al., 2015), but recent studies in a muscle-specific *Inpp5k* mouse knock-out line also determined that abnormal accumulation of PtdIns(4,5)P_2_ led to a severe disruption in lysosome recycling (McGrath et al., 2020). Interestingly, lysosome enlargement and autophagy inhibition found in the *Inpp5k*-deficient muscle were not dependent of Akt/mTOR signaling, suggesting an independent additional role for PtdIns(4,5)P_2_ in muscle maintenance in the autophagic lysosome reformation pathway (McGrath et al., 2020). In addition, increased levels of D3-phosphoglycerate dehydrogenase (PHGDH) have been found in fibroblasts obtained from individuals with INPP5K phosphatase mutations, indicating further metabolic disruptions (Hathazi et al., 2021). Overall, we propose that targeting the phosphatase domain in Inpp5ka would lead to a reliable model for *INPP5K* mutations in humans.

Whether Inpp5kb evolved to perform a different function in the pineal gland and how it lost its phosphatase activity in the zebrafish remains to be studied.

## Acknowledgements

The authors would like to thank all members of the Manzini laboratory for helpful discussion, Kathleen Flaherty and Heather Pond for management of the zebrafish colonies at Rutgers University and the George Washington University, respectively, and Himani Majumdar for assistance with the *in situ* protocols.

This work was funded by the National Institutes of Health R01NS109149 to M.C.M, Wellcome Trust (105616/Z/14/Z) and the Medical Research Council (MRC/N010035/1) to L.E.S. Additional support was provided to M.C.M. from the Robert Wood Johnson Foundation (grant #74260).

## Author contributions

M.C.M. and E.S.C. conceived the study. D.S., B.M.G., E.S.C., L.E.R., B.F.K., L.C. and L.T. performed experiments and collected data. D.S., B.M.G., E.S.C. and L.T. analyzed the data. D.S., B.M.G. and M.C.M. wrote the manuscript with contributions from S.A.M. and L.E.S. All authors reviewed the manuscript.

